# BioInformatics Agent (BIA): Unleashing the Power of Large Language Models to Reshape Bioinformatics Workflow

**DOI:** 10.1101/2024.05.22.595240

**Authors:** Qi Xin, Quyu Kong, Hongyi Ji, Yue Shen, Yuqi Liu, Yan Sun, Zhilin Zhang, Zhaorong Li, Xunlong Xia, Bing Deng, Yinqi Bai

## Abstract

Bioinformatics plays a crucial role in understanding biological phenomena, yet the exponential growth of biological data and rapid technological advancements have heightened the barriers to in-depth exploration of this domain. Thereby, we propose **B**io-**I**nformatics **A**gent (`BIA`), an intelligent agent leveraging Large Language Models (LLMs) technology, to facilitate autonomous bioinformatic analysis through natural language. The primary functionalities of `BIA` encompass extraction and processing of raw data and metadata, querying both locally deployed and public databases for information. It further undertakes the formulation of workflow designs, generates executable code, and delivers comprehensive reports. Focused on the single-cell RNA sequencing (scRNA-seq) data, this paper demonstrates `BIA`’s remarkable proficiency in information processing and analysis, as well as executing sophisticated tasks and interactions. Additionally, we analyzed failed executions from the agent and demonstrate prospective enhancement strategies including self-refinement and domain adaptation. The future outlook includes expanding `BIA`’s practical implementations across multi-omics data, to alleviating the workload burden for the bioinformatics community and empowering more profound investigations into the mysteries of life sciences. `BIA` is available at: https://github.com/biagent-dev/biagent.

## 1 Introduction

Bioinformatics is an interdisciplinary discipline, by leveraging cutting-edge computational methodologies and algorithmic strategies, it plays an irreplaceable role in many fields such as biology [1], medical science [2], microbiology [3], etc. Bioinformatics fosters a holistic understanding of biological processes by facilitating the integration and interpretation of multi-level data, from genomics to transcriptomics, proteomics, metabolomics, and beyond [4]. It helps people expand their perspectives in biological sciences and extending the limits of human cognition in the intricate tapestry of life. The progression of bioinformatics plays an indispensable role in driving the evolution of numerous fields of study.

With the continuous advancement of multi-omics sequencing technologies and the gradual reduction in costs, the output of omics data and associated analytical tools is growing at an unprecedented rate. This trend presents both significant opportunities and new challenges for researchers in the field of biological sciences. For instance, a specific workflow for the processing whole-genome sequencing (WGS) data, involves utilization of dozens of software tools ^3^ and necessitating researchers to possess fundamental skills in software installation, parameter invocation, and troubleshooting capabilities. Furthermore, the in-depth analyses demand proficiency in an expanded toolkit of coding and visualization. These prerequisites constitute a considerable hurdle for diverse stakeholders in the bio-sciences, such as the wet-lab biologists whose work are predominantly empirical, and clinicians who often lack formal training in programming [5]. Indeed, the bioinformatics communities have been actively engaged in tackling these challenges. Many projects, for instance, the Galaxy Project [6], are building low-barrier-to-use, reusable, and wide-ranging platforms that facilitate efficient storage, management, and analysis of the vast datasets for researchers. Bioconductor [7], comprises more than 2000 R packages, while the array of shell tools and Python packages has burgeoned to hundreds, all continually evolving and improving. However, these endeavors are, as yet, constrained by the limitations of preceding technologies. Presently, most solutions aimed at constructing reusable analysis pipelines rely heavily on manual labor and are inadequately adaptable, underscoring the need for more sophisticated and flexible methodologies.

Thankfully, the development of AI technologies, especially Large Language Models (LLMs) [8, 9, 10] with strong reasoning, adequate knowledge reserve and excellent coding capabilities [11], is reshaping the paradigms and precepts of how people leverage bioinformatics data. Following the introduction of ChatGPT in Nov 2022, a diverse array of LLMs has been made accessible to the public. Notable products include commercial models, such as GPT-4 from OpenAI [12] and Gemini from Google [13], and open-source alternatives, like Llama from Meta [14] and Qwen from Alibaba [15]. LLM-based agents are intelligent system built using the capabilities of LLMs. With a proper framework, agents can independently accomplish planning and execution of tasks in specific domains leveraging strategies like prompt engineering, environment awareness, reinforcement learning, and external knowledge modules [16]. Prior works have deployed and performed LLM-based tools such as `Dr BioRight` and `BioMonia` to understand the user’s needs and provide solutions through a dialog model [17].

In our experiments, we devise BioInformatics Agent(BIA), using the current state-ofthe-art language model from OpenAI, GPT-4, through the web API service (model id: `gpt-4-0125-preview`)^4^, to formulate experimental protocols grounded in provided solutions within the realm of bioinformatics. Agent is capable of executing the entire single-cell analysis pipeline, encompassing data retrieval, invocation of appropriate APIs for processing, compiles conclusions through autonomous planning, emerging as a potent facilitator in bioinformatics research. With the installation of `BIA`, we envision that bioinformatics researchers are alleviated from the burden of voluminous datasets and repetitive chores, thereby concentrate on more pivotal research questions.

## 2 Methods

`BIA` is operationalized via textual interactions with Large Language Models (LLMs) [18, 16], facilitating data extraction, analysis, and report generation through dialogues with users. `BIA` is implemented through textual prompt interaction with LLMs. Overall, the engagement with the LLM is orchestrated via four structured narrative segments [19]: the `Thought` segment instigates a reflective assessment of the task’s progression; the `Action` and `Action Input` segments direct the LLM to invoke a particular tool and specify its required inputs, thereby promoting instrumental engagement; finally, the `Observation` phase permits the LLM to interpret the result from the executed tool. Consequently, the LLM’s responses are meticulously dissected to initiate tool-driven actions or to formulate conclusive feedback addressed to the user, effectively closing the interaction loop and enhancing the system’s analytical capabilities. Domain-specific tools are imperative for augmenting the autonomy and functionality of LLM-based Agents. In this context, we present an extensive suite of tools crafted by domain specialists tailored for `BIA`. These tools are systematically classified into three categories according to their operational roles: online database management, metadata extraction, and bioinformatics workflow. The usages and implementations of the tools are described in the following sections.

### 2.1 Metadata processing and data acquisition

Firstly, tools that interact with leading online public repositories such as the European Nucleotide Archive (ENA) [20], National Center for Biotechnology Information (NCBI) [21], European Bioinformatics Institute (EBI) [22], among others, are installed, facilitating the download of DNA and RNA datasets along with their associated metadata information.

The *search online db(query)* tool provides a proxy of user queries to the search engines of the public repositories that look for relevant samples based on meta information, and returns a list of IDs of matching samples. The *download online db(ids)* tool downloads the data, including metadata, processed count matrices and raw data, from the repositories using given IDs. These tools are implemented using `ffq` [23], for fetching sequence read files and basic meta information, and `GEOparse` [24], which augments metadata of samples from GEO, in Python. In this paper, `BIA` is prompted with multiple rounds of conversations to determine a specific study and the data for it, clarifying the data needed to use only single-cell RNA data and selecting only Homo sapiens.

When acquiring count matrices for chosen samples, `BIA` accommodates a broad spectrum of data formats, including SRA, FASTQ, MTX, TSV, RData. The *read count data(ids)* tool first attempts to convert the processed count matrix uploaded by authors to a standard annotated data (Anndata) format [25]. The formats of count matrix is diverse with arbitrary column and row arrangements. Reading such data involves an understanding of the matrix structure, and thus cannot be hard-coded. We addressed this challenge via the coding skill of LLMs by prompting LLms with short summaries of uploaded files and code templates selected based on file extensions. We execute the completed replied code in a Python environment and obtain the final Anndata object. For samples without author-provided data, `BIA` builds the Anndata from the sequence read data following standard practice, i.e., aligning public FASTQ files with Cell Ranger software [26] using default settings. `BIA` then carries out data quality check, pre-processing and format conversion tasks on Anndata object.

The second critical functionality pertains to the identification and organization of metadata, which refers to structured information including patient demographics, clinical Data, and treatment and intervention for each sample [27]. Inconsistencies in metadata recording across databases hinder data reuse, while in bioinformatics, carefully control of confounding variables is vital for reliable data interpretation [28]. Here, to enrich the quality of sample metadata, we leverage LLMs to extract structured data defined by experts from unstructured raw descriptions supplied by authors. The *metadata extraction(ids)* function iterates through a list of given sample IDs and returns a predefined set of meta fields from the meta information for each sample. The meta fields, selected by experts, can act as effective filters for downstream analysis. To collect the field values, we first obtain the text descriptions of samples, i.e., the SOFT and SDRF files for samples from GEO and ArrayExpress, respectively. This raw information is then combined with instructions of target meta fields to form a textual prompt. Last, we call LLMs with the prompt and extract corresponding data for every meta fields from replies.

#### 2.1.1 Workflow tools invoker

The bioinformatics toolbox is expansive and undergoes rapid updates, featuring highly flexible workflow customization, posing a significant challenge to an agent’s capability in tool invocation and adaptation. To address this, we primarily employ the following strategies.

We obtain scRNAseq bioinformatics analysis use cases from publicly available resources (e.g., best practices, scanpy) as metadata in the input data, and describe these use cases as embedding data by human experts. These data are then cleaned and formatted as necessary to ensure the data quality. We apply OpenAI’s embedding model (model id: `text-embedding-3-small`) to convert the processed textual data into vector representations (i.e., embeddings), and construct a embedding database indexed with Chromadb package^5^. Such embeddings can be used to retrieve the most relevant use cases for a given query by finding the nearest neighbors of the query embedding [29]. We then construct a bioinformatics tool invocation wrapper as follows, while ensuring dynamism and flexibility (Figure 2).

We regulate the use of bioinformatic tools by the description of the bioinformatic task, the perception of the environment, and the user’s expectations. However, in the real-world scenario, some stable or simple workflows do not necessarily require the users’ expectations to tune the output, which we here named them static workflows, and dynamic workflows for the counterparts. Our invoker is designed to be compatible with them as shown below, we use static workflow invocation as the foundation, and add user expectation management on top of that to adapt dynamic workflows. First, *Task description* is used to query representations of the most comparable bioinformatics tool use cases reference through RAG technology. Information about variables and packages loading in the environment is abstracted into textual representation and packaged into *Environment perception*, then they are given to Agent for code usage adaptation along with the reference use case. Then the adapted code are executed by the Agent with automatic error correction. At this point the results of the static workflow have been produced, and if there is no input of the *Users’ expectations*, the invoker’s work is finished. On the other scenario, if the user’s expectations are given, the invoker will perform a dynamic workflow invocation, and the output of the static workflow will first be summarized into a linguistic representation based on the user’s needs, and then evaluated and proposal for optimization will be generated by the Agent, in which the latest resources on the network will be used to ensure that the evaluations are biologically sound. Here, we specify that the proposal are focused to the parameter adjustments of the current use case, the it is fed back to the Agent for re-adaptation of the use case until the results meet the users’ demands.

## 3 Results

Here, we present an exemplar case of comprehensive single-cell data analysis workflow, demonstrating the core functionalities of `BIA`. It primarily incorporates chat-based interaction, data retrieval upon query, automated code generation, and the subsequent aptitude for analyzing and interpreting biological datasets. The overarching framework is illustrated in Figure 1.

**Figure 1:**
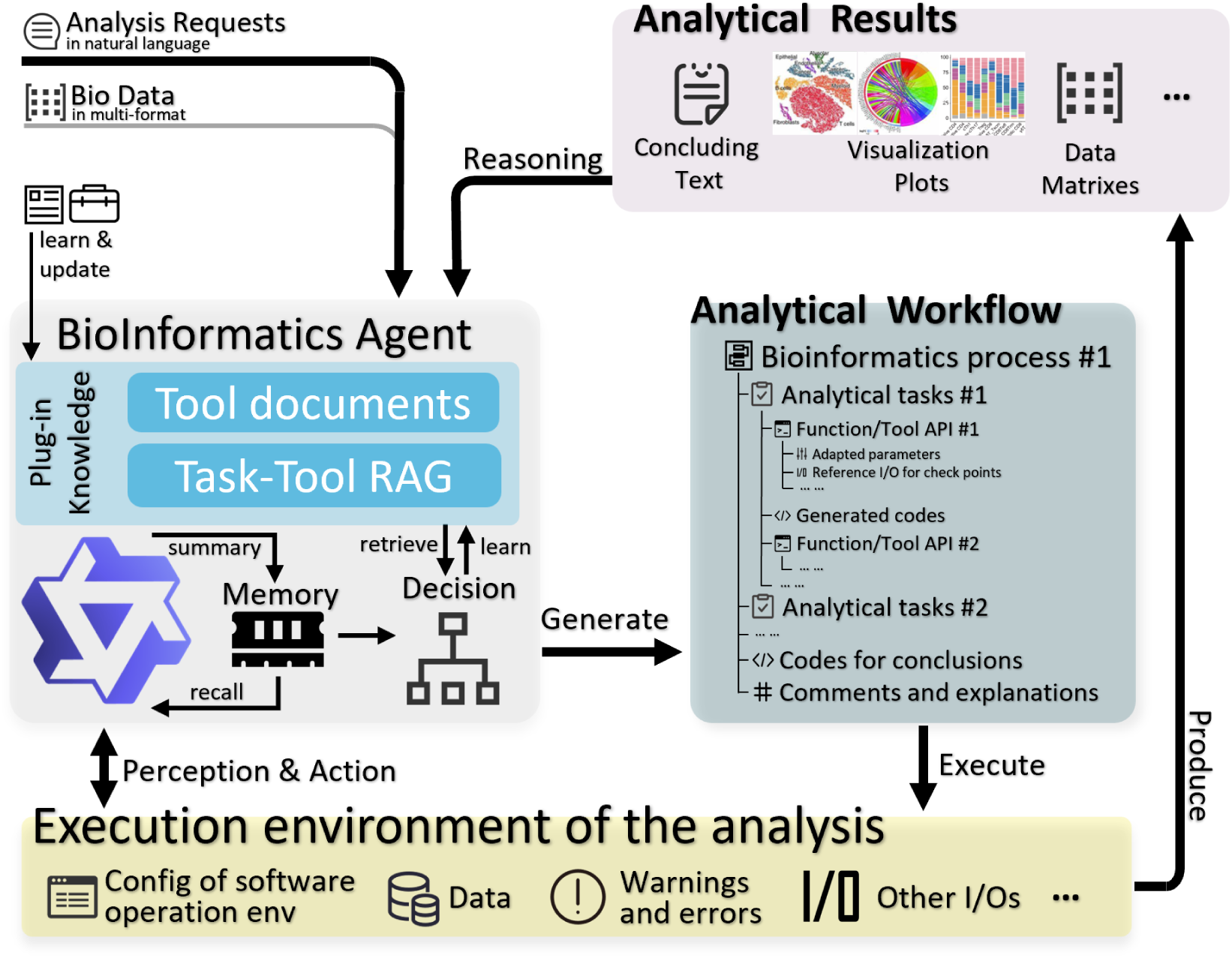
Overview of BioInformatics Agent(BIA)’s overall framework. BIA involves the following key steps: 1). Input Processing: receiving and preprocessing the user’s input or query and classifying the input into predefined categories 2). Generative Process: Given the processed input and contextual understanding, BIA deploy tools and generate workflow 3). Response Evaluation and Filtering: Generated responses are checked for coherence, appropriateness, and adherence to predefined rules. 4). Feedback Loop: Feedback used to fine-tune the model, improving its performance over time through reinforcement learning or adaptive request. 5). Delivery: Finally, the generated and processed results are delivered to the user through interface and they may use for reasoning for the model again.

**Figure 2:**
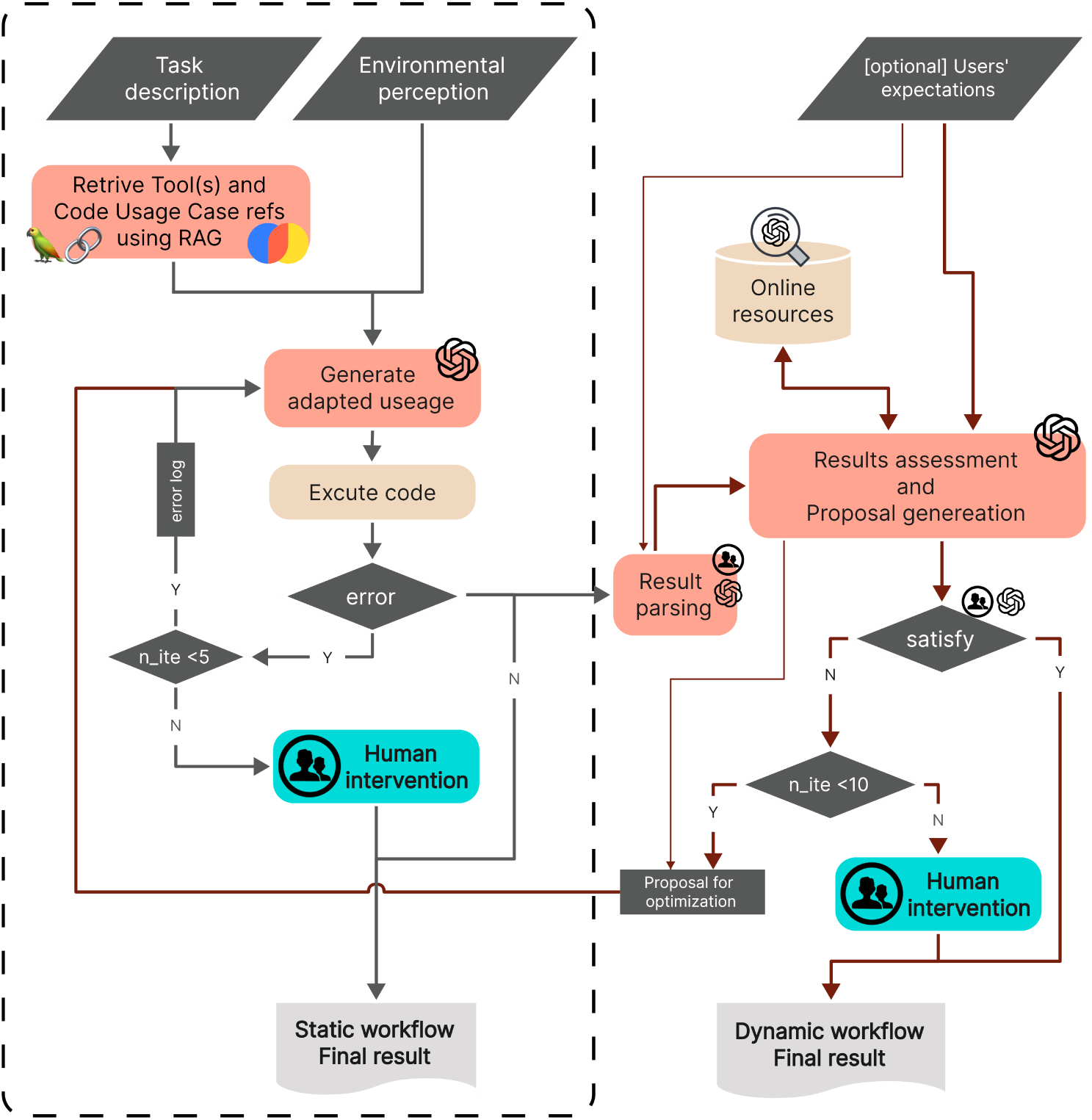
Flow chat for informatics tool invoker. The user initiates the process by providing a detailed task description along with optional anticipated computational outcomes within the invoker. Leveraging LLMs, the system mines the RAG database in response to the task description, extracting pertinent toolkits, code snippets, and usage scenarios. Integrating the user’s specifications with environmental insights garnered by the agent, the retrieved references are synthesized to regenerate contextually adapted tool utilization code. The tailored code then undergoes invocation, execution, and iterative tweaking. The process is dynamically guided, ensuring alignment with evolving requirements. Final step, the resultant outputs are produced, and when provided, the agent benchmarks these against the user’s predefined expectations for computational outcomes. In instances where user expectations are specified, the agent actively engages in a feedback cycle, tapping into online resources to evaluate and, if necessary, rectify the output. Human intervention is solicited at critical junctures to ensure accuracy and relevance.

### 3.1 Automated scRNA data collection

`BIA` offers an interface enabling users to query content from a locally deployed database, as well as to download openly accessible datasets. The first experiment we designed a user’s request : “Find single-cell transcriptomic studies involving pancreatic cancer from local database.” Based on this requirement, `BIA` will first check local database for relevant samples and return a list of IDs of matching samples along with sorted metadata, processed count matrices and raw data, from the repositories using given information as show in Figure 3.

**Figure 3:**
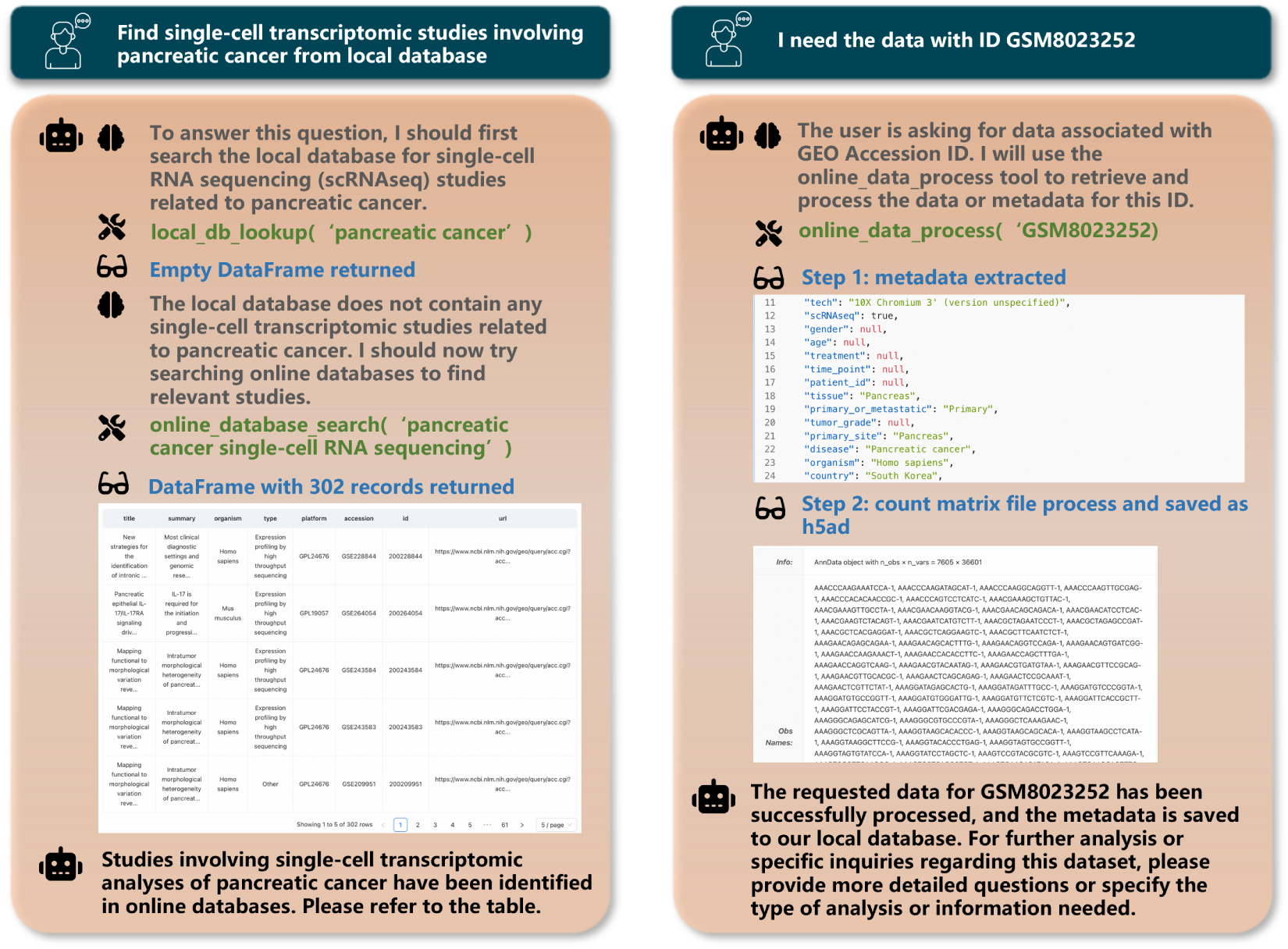
BIA searches datasets from local and public databases. A user case of the `BIA` pulls dataset and return with results.

If the local database query return an empty result for pancreatic cancer samples, the system initiates a call to the ffq software to retrieve dataset URLs and accompanying text descriptions for each Run/Sample from public databases, such as GEO or ArrayExpress. Thereafter, descriptive metadata is consolidated and formatted as depicted in Figure 3, facilitating downstream analyses. Matrix data is either directly downloaded or processed from raw data into mtx format utilizing tools like cellranger. These steps have been relatively fixed through engineering, and the `BIA` plays a major role in calling the API and executing it (Figure 1).

### 3.2 Experimental design generation

In the following experiments, we tested the ability of `BIA` to intelligently invoke and automatically adapt the bioinformatics analysis process in a code-free interaction. We selected 2 datasets containing single-cell RNA-seq data of triple-negative breast cancer(TNBC) samples from the GEO database (GSE143423, GSE148673) and selected 6 of the samples (GSM4259357, GSM4476486– GSM4476490). Then, according to the task description entered by the natural language, the `BIA` can call the bioinformatics analysis tool via Tool-invoker to complete the user-specified task.

In the course of executing bioinformatics analyses, we categorize bioinformatics analysis workflows into static procedures and dynamic procedures, depicted respectively as yellow and red in Figure 4, based on the relative maturity and degree of personalization of the bioinformatics tasks. Static procedures resemble automated script executions, wherein the `BIA` invokes reference code from the retrieval augmented generation (RAG) [29] according to the task description, adjusts the code and parameters in accordance with specific parameters and task details, and subsequently executes the code to carry out the bioinformatics analysis, yielding analytical results. On the other hand, dynamic procedures embody AI-augmented personalized analysis. Building upon fixed procedures, we incorporate an additional step enabling the `BIA` to iteratively refine the analysis script. This involves the `BIA` first summarizing and analyzing the output from the initial run, comparing these results against the user’s anticipated outcomes. Should the determination be that the results do not align with user expectations, the `BIA` then modifies the entire workflow to ensure the user’s objectives are met. This iterative process of self-refinement within the adaptive workflow represents a significant leap in tailoring bioinformatics analyses to individual researcher needs and enhancing overall analytical flexibility.

**Figure 4:**
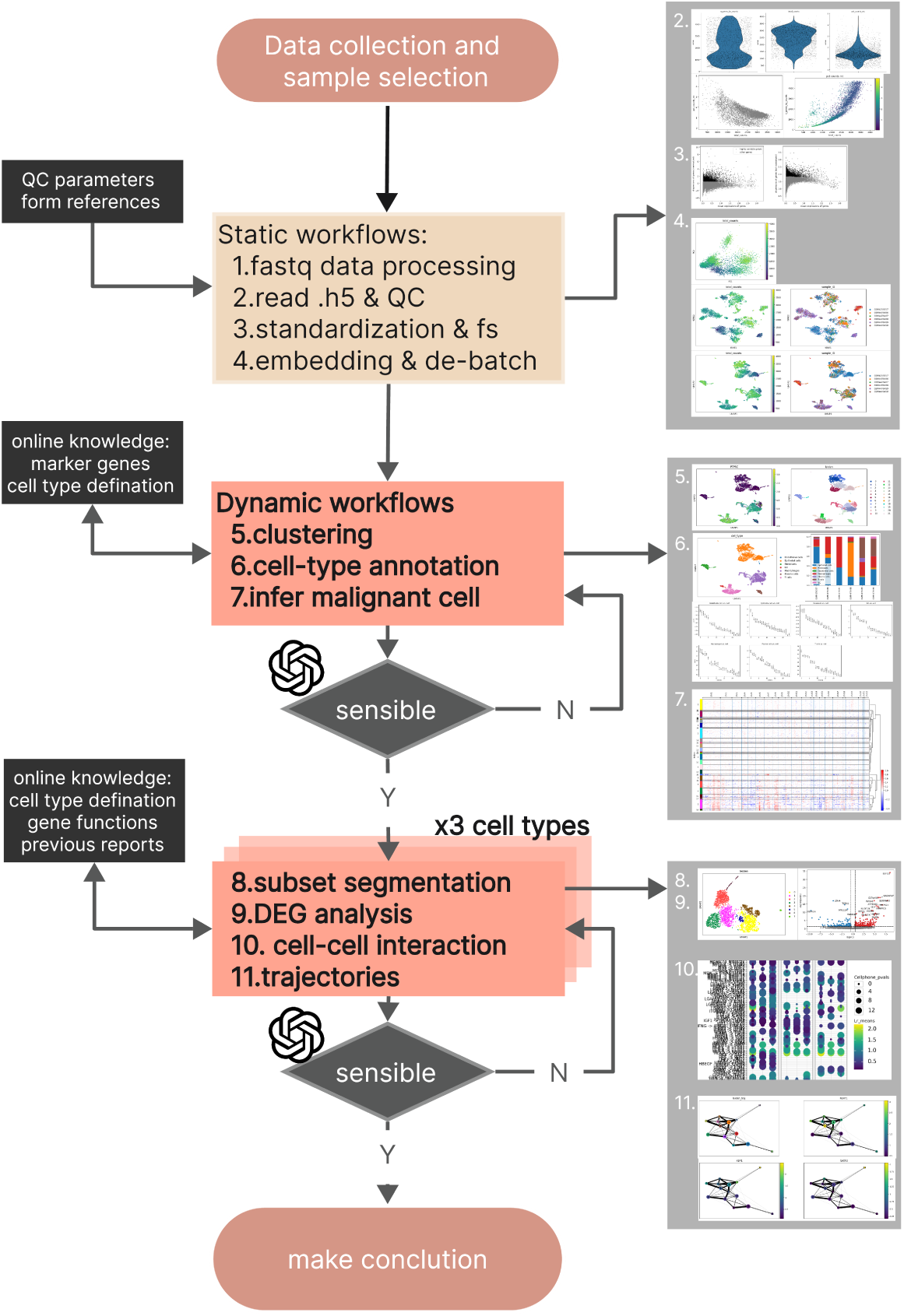
Bioinformatics analysis process and results overview by BIA. The diagram presents BIA completes a comprehensive bioinformatics analysis framework through natural language instructions. 6 TNBC sampels were selected form 2 datasets. It encompasses 11 scRNA-seq analysis tasks, and the results are placed in the right.

We wrap 11 commonly used bioinformatic analysis tasks/processes (Figure 4) and experimented with performing bioinformatic analysis with `BIA` using a code-free approach. In this experiment, we completed the testing of 11 bioinformatic analysis tasks, including data reading and quality control, data normalization and feature screening, data dimensionality reduction mapping and de-batching, cell clustering and subgrouping, cell type annotation, malignant cell inference, cell subpopulation segmentation, differential gene analysis, cell interaction analysis, and pseudo trajectory analysis. For static workflows, BIA efficiently completes the tasks within three iterations. However, when undertaking dynamic workflows, such as subset segmentation, manual intervention tends to be indispensable to guarantee precision and tailor the process to particular needs.

### 3.3 Advanced analytics and reporting

`BIA` is capable of invoking data outcomes from preceding steps based on user requirements, as well as retrieving previously stored results from the database, to compile and present comprehensive analytical reports. It can perform sophisticated statistical analyses like compare different clusters as required to make data-driven decisions and predictions with greater accuracy. The examples were presented in Figure 5. In a case study, we posed the query: Based on the data in our local database, show me the level of CD4+ T cells in the breast cancer sample with id “LUAD-003-01-1A”. `BIA` will return the proportion of each cell type like precisely delivering the composition of distinct CD4+ T cell subsets in response to the input request. Further leveraging its LLM knowledge base, BIA is capable of identifying the common markers for each cell type and furnishing encyclopedic explanations rooted in domain expertise. This significantly reduced the cognitive load and time investment for users in querying and interpreting the results, thereby enhancing the efficiency and accessibility of bioinformatics data analysis.

**Figure 5:**
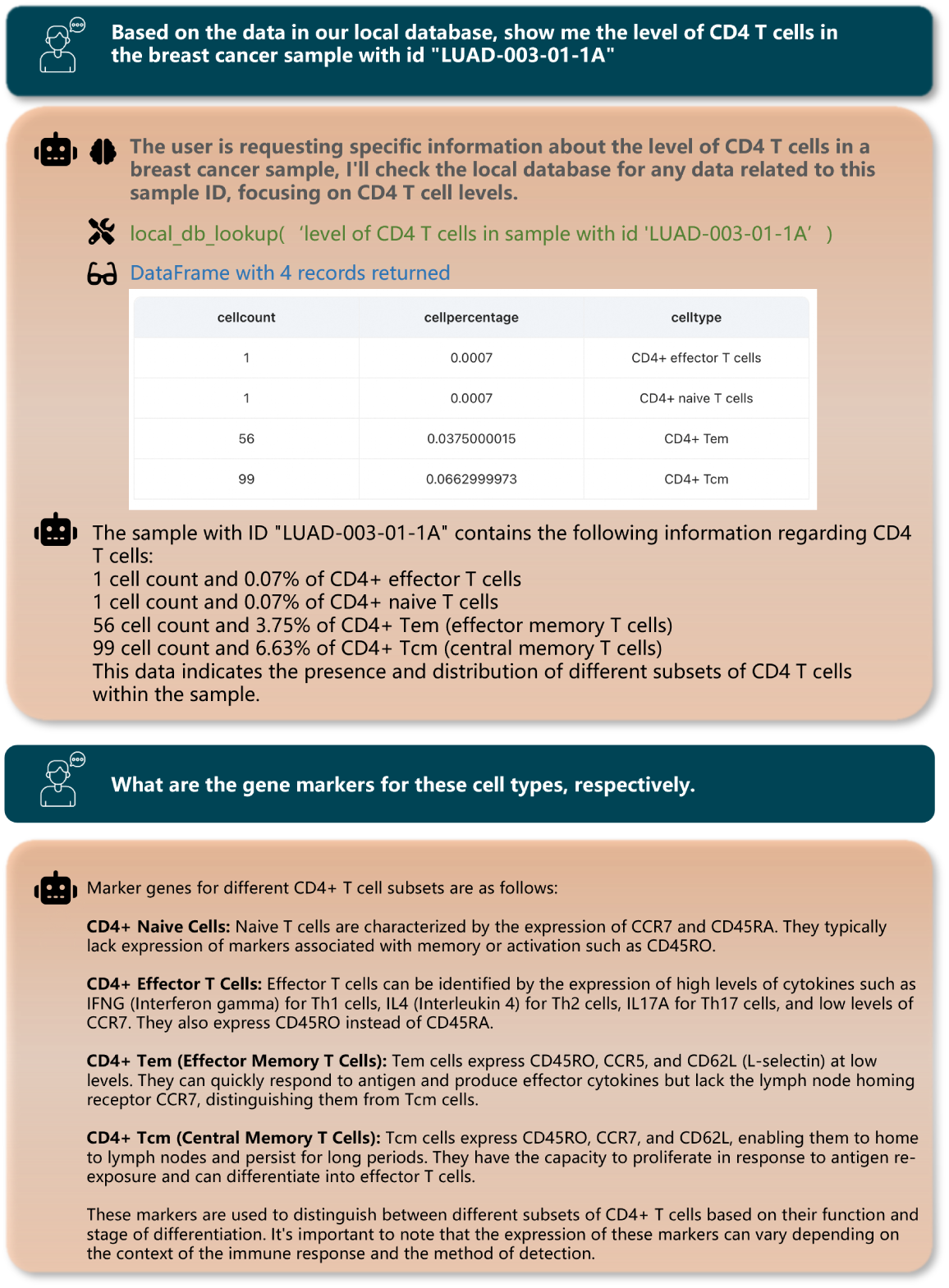
Advance reporting demo case. The `BIA` queries and characterises database results through natural language and expands conclusions based on online knowledge.

### 3.4 Optimistic outcome and strenuous challenges

Developing a fully tailored LLM and toolsets specifically for the realm of biology presents a formidable challenge. Through thoroughly evaluation of the `BIA`s’ performance across each stage of tasks, we have garnered promising outcomes while simultaneously uncovering several common issues, the following section describe the evaluation results of the `BIA`, common issues and their causes, and the direction of subsequent adjustments.

We first assessed the effectiveness of metadata extraction accuracy of `BIA`. We obtained over 3k textual metadata representations from the GEO (NCBI) and Array Express (ENA) databases, and had the `BIA` performed 11 metadata categories (including gender, age, disease, sampling site and other information) extraction from them (Section 2). It achieved more than 95% accuracy in both GEO and ArrayExpress metadata extractions, and in very irregular dataset (GEO Hard, for which the authors of this section provide very little information), it meets expectations in most factors. We will publish the 3k Meta Ground Truth dataset along with the paper.

Subsequently, we examined the `BIA`’s capability in designing bioinformatics experiments, encompassing both static and dynamic processes. In the context of fixed procedures, adhering to established pipelines for single-cell analysis, `BIA` was capable of invoking pre-packaged codes that could be readily modified with parameters tailored to our specific requirements, thus aligning with predefined expectations. In contrast, during the zero-shot dynamic processes, we observed certain limitations. The resultant experimental designs for bioinformatics analyses were found wanting in completeness; crucial steps such as sub-cluster annotation and trajectory analysis were occasionally omitted. Moreover, there was a notable inconsistency in the results, with varying tools being suggested as missing from the experimental protocol when confronted with identical queries, indicative of a lack of stability and robustness in the adaptive process. Other analogous issues encountered with large-scale language models encompass instances where generated code snippets either fail to fulfill their intended functionality or prove altogether non-executable. These issues can be attributed to several underlying factors: the LLM’s dearth of domain-specific knowledge; the propensity for “systematic hallucinations” during model processing; a heavy reliance on short-term memory mechanisms; and a relatively low degree of internal logical consistency. Additionally, the inability of such models to autonomously iterate through decision-making processes, necessitating manual intervention to define loop termination, may stem from inadequate information provisioning or insufficient automated evaluation mechanisms.

In summary, the `BIA` has demonstrated a remarkable level of autonomy and sophisticated biological awareness in conducting bioinformatics research. However, further enhancements are required to optimize its performance. Key directions for future work include enriching the Agent with more specialized knowledge to elevate the sophistication of experimental designs it generates, developing a well-aligned prior knowledge base and integrating it into the Agent via fine-tuning or long-term memory mechanisms, enhancing the Agent’s ability to interpret and generate structured data, and refining its contextual understanding and reasoning capabilities. These experiences will be applied to additional omics datasets with the aim of significantly improving the LLM’s proficiency in bioinformatics.

## 4 Discussion and Conclusion

Herein, we introduce the `BIA`, a system adept at substantially augmenting the efficacy of bioinformatics analyses while concurrently diminishing the entry barriers to the biological domain knowledge. This paper primarily on a single-cell RNA sequencing (scRNA-seq) analysis case, delineates a workflow commencing with data acquisition, proceeding through bioinformatic experiment design and execution, and ultimately concluding with the autonomous organization and interpretation of analytical outcomes by the Agent. This end-to-end process underscores the capacity of the `BIA` to streamline complex bioinformatics tasks, thereby enhancing productivity and deepening biological insights. Notably, the `BIA` excels in converting arbitrary linguistic descriptions into structured metadata, which greatly enhancing the exploratory power of large cohort datasets, and significantly boosting bioinformatics productivity. Nevertheless, we have also identified a number of issues that need to be addressed, such as low self-consistency when planning and hallucinations. To address these, our future refinements encompass fine-tuning for domain specificity and reinforcement learning augmented with human feedback for iterative performance enhancement.

Bioinformatics is evolving towards being ever more burdensome and a rising threshold. We aspire to enhance the generalizability of bioinformatics methodologies to mitigate the constraints imposed by both human capital and computational resources. It is imperative to acknowledge that the `BIA`, while promising, upon substantial refinements, rigorous validation and a focused enhancement of its intelligence and accessibility,it can universally adopted in the broader biological sciences community.

## Footnotes

3 https://gatk.broadinslitute.org/hc/en-us/sections/360007226651-Best-Practices-Workflows

4 https://platform.openai.com/

5 https://github.com/chroma-core/chroma

